# Time-delayed biodiversity feedbacks and the sustainability of social-ecological systems

**DOI:** 10.1101/112730

**Authors:** A.-S. Lafuite, M. Loreau

## Abstract

The sustainability of coupled social-ecological systems (SESs) hinges on their long-term ecological dynamics. Land conversion generates extinction and functioning debts, i.e. a time-delayed loss of species and associated ecosystem services. Sustainability theory, however, has not so far considered the long-term consequences of these ecological debts on SESs. We investigate this question using a dynamical model that couples human demography, technological change and biodiversity. Human population growth drives land conversion, which in turn reduces biodiversity-dependent ecosystem services to agricultural production (ecological feedback). Technological change brings about a demographic transition leading to a population equilibrium. When the ecological feedback is delayed in time, some SESs experience population overshoots followed by large reductions in biodiversity, human population size and well-being, which we call environmental crises. Using a sustainability criterion that captures the vulnerability of an SES to such crises, we show that some of the characteristics common to modern SESs (e.g. high production efficiency and labor intensity, concave-down ecological relationships) are detrimental to their long-term sustainability. Maintaining sustainability thus requires strong counteracting forces, such as the demographic transition and land-use management. To this end, we provide integrative sustainability thresholds for land conversion, biodiversity loss and human population size - each threshold being related to the others through the economic, technological, demographic and ecological parameters of the SES. Numerical simulations show that remaining within these sustainable boundaries prevents environmental crises from occurring. By capturing the long-term ecological and socioeconomic drivers of SESs, our theoretical approach proposes a new way to define integrative conservation objectives that ensure the long-term sustainability of our planet.

## 1. Introduction

Current trends in human population growth (Cohen, 2003; Gerland et al., 2014) and environmental degradation (Vitousek et al., 1997) raise concerns about the long-term sustainability of modern societies, i.e. their capacity to meet their needs “without compromising the ability of future generations to meet their own needs” (Brundtland et al., 1987). Many of the ecosystem services supporting human systems are underpinned by biodiversity (Cardinale et al., 2012), and current species extinction rates threaten the Earth’s capacity to keep providing these essential services in the long run (Pereira et al., 2010; Ehrlich and Ehrlich, 2013). The long-term ecological feedback of ecosystem services on human societies has been largely ignored by neo-classical economic theory, mainly due to the focus on short-term feedbacks, and the assumption that ecosystem processes can be substituted for by human capital (e.g. tools and knowledge) and labor, thereby releasing ecological checks on human population and economic growth (Boserup, 1965; Dasgupta and Heal, 1974). In particular, tremendous increases in agricultural productivity resulted in more than 100% rise in aggregate food supply over the last century (Schmidhuber and Tubiello, 2007).

However, the substitution of human capital for natural resources, also called the “technology treadmill” (Pezzey, 1992), is currently facing important limitations. One such limitation is land scarcity, as the remaining arable land reserve might be exhausted by 2050 (Lambin and Meyfroidt, 2011). Moreover, recent projections suggest a slowdown in the growth rate of agricultural Total Factor Productivity (TFP), which measures the effect of technological inputs on total output growth relative other inputs (Kumar et al., 2008). Technological improvements may not compensate for arable land scarcity (Zeigler and Steensland, 2015), thus questioning the potential for continued TFP growth in the future (Gordon, 2012; Shackleton, 2013). Another limitation comes from the loss of many biodiversity-dependent ecosystem services which play a direct or indirect role in agricultural production, such as soil formation (Barrios, 2007), nutrient cycling, water retention, biological control of pests (Martin et al., 2013) and crop pollination (Gallai et al., 2009; Garibaldi et al., 2011). Substituting ecological processes with energy and agrochemicals has mixed environmental impacts, with unintended consequences such as water use disturbance, soil degradation, and chemical runoff. These effects are responsible for a slowdown in agricultural yield growth since the mid-1980s (Pingali, 2012) and have adverse consequences on biodiversity, human health (Foley et al., 2005) and the stability of ecosystem processes (Loreau and de Mazancourt, 2013).

Moreover, biodiversity does not respond instantaneously to land-use changes. Habitat fragmentation (Haddad et al., 2015) increases the relaxation time of population dynamics (Ovaskainen and Hanski, 2002) - i.e. how fast a species responds to environmental degradation. As a consequence, species extinctions are delayed in time, which generates an extinction debt (Tilman et al., 1994) and a biodiversity-dependent ecosystem service debt (Isbell et al., 2015). These ecological debts may persist for more than a century, and increase as species get closer to extinction (Hanski and Ovaskainen, 2002). The accumulation of these ecological debts may have long-term effects on modern human societies.

Such time-delayed ecological feedbacks have been neglected by the most influential population projection models (Meadows et al., 1972, Meadows et al., 2004; Turner, 2014). Yet, environmental degradation can have catastrophic consequences for human societies even without any delayed effect (Diamond, 2005; Ponting, 1991). A well-known example is Easter Island, this Polynesian island in which civilization collapsed during the 18th century due to overpopulation, extensive deforestation and overexploitation of its natural resources (Brander, 2007). In order to investigate the mechanisms behind this collapse, Brander and Taylor (1998) modeled the growth of the human population as endogenously driven by the availability of natural resources, the depletion of which was governed by economic constraints. Their model was essentially a Lotka-Volterra predator-prey model, which is familiar to ecologists, with an economic interpretation. It showed that one of Easter Island’s ecological characteristics - the particularly low renewal rate of its forests - may be responsible for the famine cycles which brought forward the collapse of this civilization.

Given the unprecedented rates of current biodiversity and ecosystem service loss (Pereira et al., 2010), accounting for their long-term feedback on modern human societies appears crucial. In an attempt to delimit safe thresholds for humanity, a 10% loss in local biodiversity was defined as one of the core ‘planetary boundaries^11^ which, once transgressed, might drive the Earth system into a new, less desirable state for humans (Scholes and Biggs, 2005; Steffen et al., 2015). However, in practice, the biodiversity threshold above which ecosystem processes are significantly affected varies among ecosystems (Hooper et al., 2012; Mace et al., 2014), and is correlated to other thresholds such as land-use change (Oliver, 2016). The definition of integrative thresholds thus requires that we consider the interaction between the economic and ecological components of systems.

In order to explore the long-term consequences of ecological debts for human societies, we build upon Brander and Taylor’s framework, but we allow the human population to produce its own resources through land conversion. In our approach, terrestrial natural habitats provide essential ecosystem services to the agricultural lands - which are assumed to be unsuitable to biodiversity. Biodiversity and human population dynamics are coupled through the time-delayed effect of biodiversity-dependent ecosystem services on agricultural production (ecological feedback). It may be viewed as a minimal social-ecological system (SES) model that couples basic insights from market economics and spatial ecology. From market economics, we derive human *per capita* consumptions and the rate of land conversion. From spatial ecology, we use a classical species-area relationship (SAR) to capture the dependence of species diversity on the remaining area of natural habitat, and account for ecological debts through the relaxation rate of communities following habitat loss.

We investigate the behavior of the system at equilibrium analytically, and then numerically evaluate the trajectories to the equilibrium. We show that the transient dynamics of an SES depends on its ecological, economic, demographic and technological parameters. Some SESs experience large population overshoots followed by reductions in biodiversity, human population size and well-being - which we call ‘‘environmental crises”. We then analytically derive an integrative sustainability criterion that captures the vulnerability of an SES to such crises. This criterion allows assessing the effects of some parameters on the long-term sustainability of the SES, and deriving integrative land conversion and biodiversity thresholds.

## 2. Methods

### 2.1. Coupling human and ecological dynamics

We model the long-term dynamics of three variables: the human population (H), technological efficiency (T) and biodiversity (B).

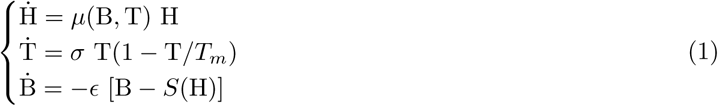

The human population endogenously grows at a rate μ(B, T), which is explicitly defined as a function of the *per capita* agricultural and industrial consumptions in section 2.1.1. Technological efficiency is assumed to follow logistic growth at an exogenous rate *σ,* until a maximum efficiency *T_m_* is reached (section 2.1.2). We use an economic general equilibrium framework to derive *per capita* human consumptions at market equilibrium, i.e. when production supply equals the demand of the human population (section 2.1.3). Using these consumptions, we derive a proportional relationship between land conversion and the size of the human population (section 2.1.4). Land conversion affects biodiversity through a change in the long-term species richness *S*(*H*). Current biodiversity B reaches its long-term level *S*(*H*) at a relaxation rate ϵ (section 2.2). Biodiversity-dependent ecosystem services then feed back on agricultural production and affect the *per capita* agricultural consumption and the human growth rate, *μ*(B, T). Model structure is summarized in Fig. 1.

**Figure 1:**
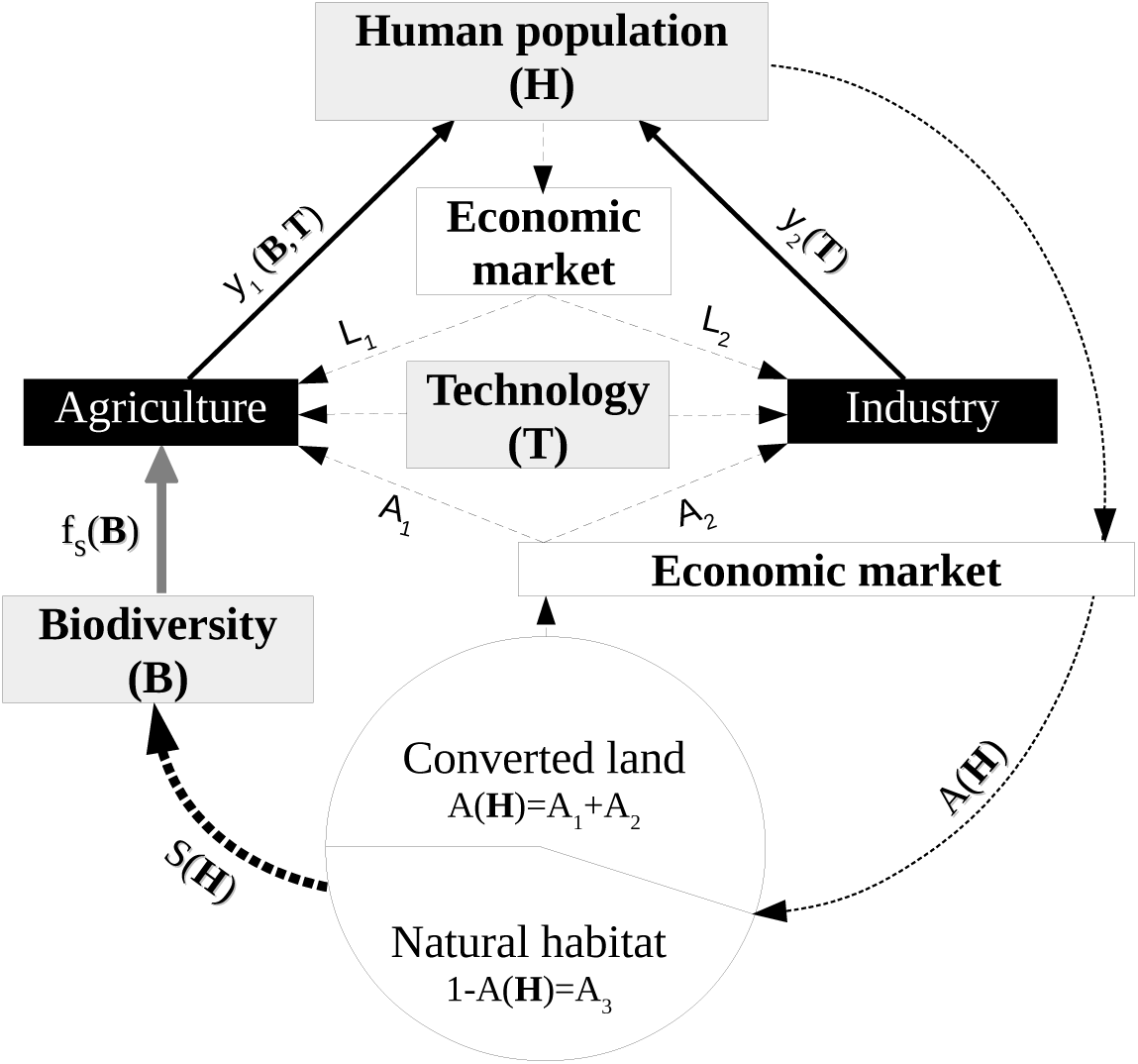
Coupling between human and ecological dynamics. Black boxes: production sectors; grey boxes: dynamical variables; white box: auxiliary economic model; dashed lines: production inputs (labor *L_i_*, land *A_i_* and technology *T*); solid lines: *per capita* outputs (*y_i_*); grey line: biodiversity-dependent ecosystem services (ecological feedback, *f_s_*); black dotted line: effect of land conversion on biodiversity; circle: total land divided into converted land (*A*_1_ + *A*_2_) and natural habitat (*A*_3_), where *A*_1_ + *A*_2_+ *A*_3_ = 1.

#### 2.1.1. Human demography

The interaction between human population, technology and income has been mainly studied by endogenous growth theory, which distinguishes three phases of economic development (Galor and Weil, 2000; Kogel and Prskawetz, 2001): (1) a Malthusian regime with low rates of technological change and high rates of population growth preventing *per capita* income to rise; (2) a Post-Malthusian Regime, where technological progress rises and allows both population and income to grow, and (3) a Modern Growth regime characterized by reduced population growth and sustained income growth (Peretto and Valente, 2015). Transition to this third regime results from a demographic transition which reverses the positive relationship between income and population growth.

In order to consider the basic linkages between human demography and economics, the growth rate of the human population is assumed to depend on the *per capita* consumptions of agricultural *y*_1_ and industrial goods *y*_2_ (Anderies, 2003):

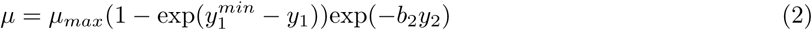

where *μ_max_* is the maximum human population growth rate, and 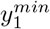 is the minimum *per capita* agricultural goods consumption, such that human population size increases if 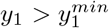, and decreases if 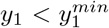 is the sensitivity of *μ* to industrial goods consumption, and thus captures the strength of the demographic transition. A higher agricultural goods consumption increases the net human growth rate while a higher industrial goods consumption eventually limits the net growth rate of the human population. Note that both *y*_1_ and *y*_2_ vary with the states of the system (section 2.1.3). The system reaches its long-term equilibrium 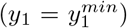 when further technological change and habitat conversion no longer increase total agricultural production, i.e. no longer compensate for the negative ecological feedback on agricultural production.

#### 2.1.2. Technological change

Technological change is central to explain the transition from a Malthusian to a modern human population growth regime. Technology is often captured through the Total Factor Productivity term (TFP) of a production function, which accounts for effects of technological inputs on total output growth relative to the other inputs, i.e. labor and land in our model. Accelerating TFP growth in recent years partially compensated for the slowing down in input growth (especially land) and allowed total output growth to maintain itself around 2% per year (Fuglie, 2008). However, recent reviews suggest that agricultural TFP growth is slowing down in a number of countries (Kumar et al., 2008) and that this trend is likely to continue (Gordon, 2012; Shackleton, 2013).

In our model, technological change increases production efficiency in the agricultural and industrial sectors, leading to higher productions for a given level of inputs (i.e. converted land and labor). We assume a logistic growth of production efficiency towards a maximum efficiency denoted by *T_m_*, where *σ* is the exogenous rate of technological change (system (1)). We explore other forms of technological change in Appendix A.1.

#### 2.1.3. Instantaneous market equilibrium

We assume a closed market where total labor is given by the size of the human population. Consumption and production levels are derived by solving for a general market equilibrium, where prices of the production inputs and consumption goods vary endogenously. As the economic dynamics are much faster than the demographic and ecological dynamics, the market is assumed to reach an equilibrium between supply and demand instantaneously.

On the demand side, the human population is assumed to be homogeneous, i.e. composed of H identical agents. *Per capita* consumptions of agricultural and industrial goods (*y*_1_ and *y*_2_, resp.) are derived from the maximization of a utility function, *U*(*y*_1_,*y*_2_), which is a common economic measure of the satisfaction experienced by consumers (Appendix A.2.1):

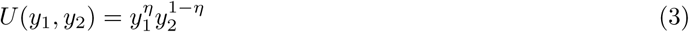

where *y*_1_ is *per capita* agricultural demand, *y*_2_ is *per capita* industrial demand, and *η* is the preference for agricultural goods.

On the supply side, a total quantity *Y*_i_ of goods is produced by sector *i,* using two production inputs, labor *L_i_,* and converted land *A_i_* (Appendix A.2.2):

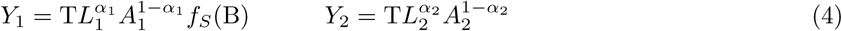

where production efficiency is captured by the variable T, *α_i_* is the relative use of labor compared to land in sector i, and the function *f_s_*(B) measures provisioning of biodiversity-dependent ecosystem services to agricultural production (ecological feedback). Such Cobb-Douglas production functions with constant returns to scale are a common assumption of growth models and allow for the substitution of land by labor. Potential ecological feedbacks on industrial production were ignored.

When demand equals supply, the *per capita* consumptions are (Appendix A.2.3):

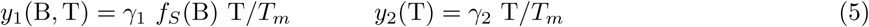

where γ_1_ and γ_2_ are explicitly defined as functions of the parameters of the system in Table 1.

**Table 1:**
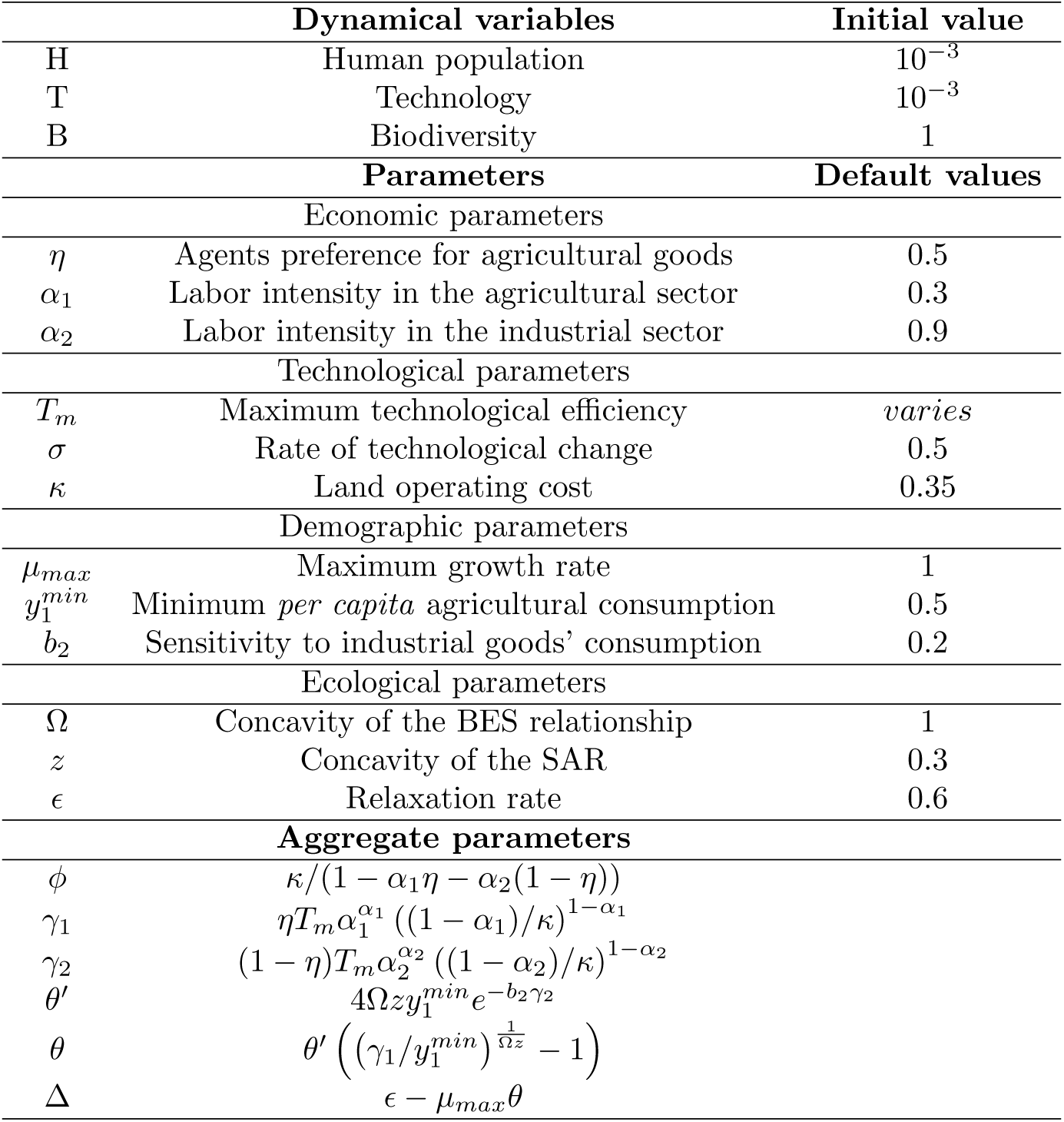
Definition and default values of the parameters and dynamical variables.

The *per capita* consumption of agricultural goods y1 is thus subject to the trade-off between the negative effect of decreasing biodiversity-dependent ecosystem services, and the positive effect of technological change. Conversely, the *per capita* consumption of industrial goods *y_2_* monotonously increases with technological efficiency - as does the strength of the demographic transition.

#### 2.1.4. Land conversion dynamics

Rising agricultural and industrial productions require the conversion and maintenance of land surfaces *A*_1_ and *A*_2_ respectively, at a cost of *κ* units of labor per unit of converted area. Land conversion reduces the area of natural habitat *A*_3_, where *A*_1_ + *A*_2_ + *A*_3_ = 1. Let us denote by A the converted land, i.e. *A* = *A*_1_ + *A*_2_. At the market equilibrium, the relationship between *A* and the human population size H is (Appendix A.2.4):

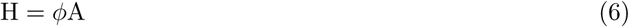

where *ϕ* represents the density of the human population on converted land, and is explicitly defined as a function of the economic parameters in Table 1. The converted area A decreases with the land operating cost *κ*, since high operating costs reduce the incentive to convert natural habitat. Proportional relationships between human population sizes and converted surfaces are commonly observed in local and regional developing economies, where subsistence agriculture remains strong and the transition to a modern growth regime is not achieved yet (Meyer and Turner, 1992).

### 2.2. Spatio-temporal dynamics of biodiversity and ecosystem services

We need two types of relationships, which capture (1) the dependence of biodiversity upon the area of natural habitat *A*_3_ and the size of the human population (*S*(*H*)), and (2) the dependence of ecosystem services upon biodiversity (*f_s_*(*B*)). Since species richness can be related to the area and spatial characteristics of natural habitats (Mac Arthur and Wilson, 1967), we choose species richness as a biodiversity measure in our system.

#### 2.2.1. Species-area relationship (SAR)

SARs are commonly used tools in ecological conservation to assess the effect of natural habitat destruction on species richness (Pereira et al., 2014). A common way to describe the decrease of species richness with habitat loss is to use a power function 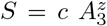, where *c* and *z* represent the intercept and the slope in log-log scale. By choosing a unitary intercept (*c* = 1) and under the constraint *A*_1_ + *A*_2_ + *A*_3_ = 1, the number of species *S*(*H*) that can be supported in the long run on an area *A*_3_ of natural habitat can be rewritten:

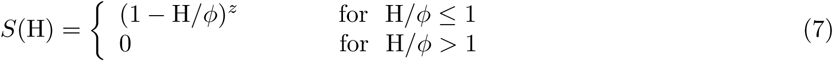

In terrestrial systems, the slope *z* typically ranges from 0.1 to 0.4 (Connor and McCoy, 1979; McGuiness, 1984), with values depending on ecosystem characteristics and species response to habitat loss. In our model, the long-term species richness *S*(H) thus decreases when the human population grows, and equals zero if the human population exceeds its maximum viable size *ϕ*.

We assume, following empirical (Ferraz et al., 2007; Wearn et al., 2012) and theoretical (Mac Arthur and Wilson, 1967; Diamond, 1972) expectations, that the rate of community relaxation to this long-term richness is proportional to the difference between current richness, *B,* and long-term richness *S*(*H*) (system (1)), where the relaxation rate is *ϵ* (Diamond, 1972).

#### 2.2.2. Biodiversity-ecosystem service relationship (BES)

Species richness has a positive effect on the level of many regulating services (Cardinale et al., 2012), among which the pollination and pest regulation services are particularly important to agricultural production - since the production of over 75% of the world’s most important crops and 35% of the food produced is dependent upon animal pollination (Klein et al., 2007).

The relationship between biodiversity and ecosystem services can be captured by a power function (Kremen, 2005; O’Connor et al., 2017) :

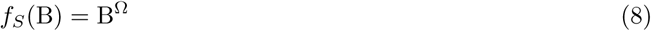

where Ω < 1, since the shape of the function *f_s_* is mostly concave-down in terrestrial systems (Mora et al., 2014).

### 2.3. Model summary

System (1) can be rewritten as:

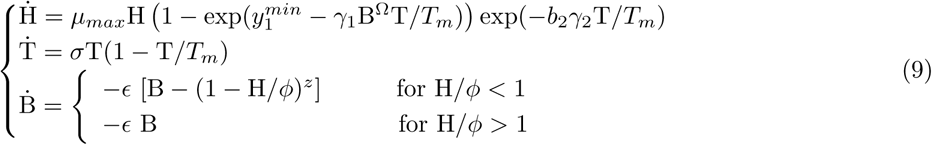

The aggregate parameters γ_1_, γ_2_, and *ϕ* result from the economic derivations of sections 2.1.3, 2.1.4 and Appendix A.2. All parameters and aggregate parameters are explicitly defined in Table 1. In the short term, technological change and human population growth drive land conversion and increase human consumption and well-being. In the long term, the time-delayed biodiversity loss reduces the supply of biodiversity-dependent ecosystem services to agricultural production (ecological feedback). The ecological relaxation rate *ϵ* generates a time lag between human and ecological dynamics, mediated through the ecological feedback. In the next section, we investigate the consequences of this time lag on the long-term sustainability of an SES.

## 3. Results and Discussion

### 3.1. Analysis of the dynamical system

For a given set of parameters and initial conditions, the dynamics of our SES is driven by the interaction between the dynamical variables summarized in eq. (9). In section 3.1.1, we characterize the steady states that can potentially be reached by the SES in the long term, and the necessary condition for their stability. We then simulate in section 3.1.2 the transient dynamics of the SES over time, for various time lags between the ecological and human dynamics (*ϵ*).

#### 3.1.1. Steady states and stability condition

A steady state is reached by the dynamical variables when the system is at equilibrium, i.e. when Ḣ = Ṫ = Ḃ = 0 (Appendix A.3.1). There are two potential steady states (*H,T,B*) for our system: (1) an undesirable equilibrium (0, *T_m_*, 1), when the parameters do not allow the human population to persist in the long term, and (2) a desirable equilibrium (*H^*^,T_m_, B^*^*):

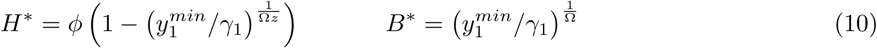

For a given set of parameters, only one of these equilibria is stable and reached by the SES (Appendix A.3.2). The desirable equilibrium (*H^*^,T_m_, B^*^*) is stable if condition (11) is met:

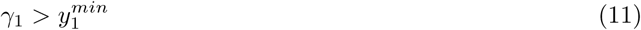

Condition (11) captures the capacity of an SES to persist in the long term, given its technological and economic characteristics (Table 1). The higher Yi, the higher the initial human growth rate and the lower the biodiversity at equilibrium B*. When condition (11) is (not) met, the SES reaches the desirable (undesirable) equilibrium in the long term. According to eq. (10), natural habitat (1 — H/*ϕ*) is never entirely converted at equilibrium, since *H*/*ϕ* < 1.

Condition (11) only guarantees that the desirable equilibrium is reached in the long term, but tells nothing about the trajectories to the equilibrium, and especially about the effect of the relaxation rate *ϵ*.

#### 3.1.2. Transient dynamics for varying relaxation rates *ϵ*

In order to test for the effect of a time-delayed ecological feedback on the transient dynamics of an SES, we numerically evaluate the trajectories to the equilibrium and increase the lag between the human and ecological dynamics by decreasing *ϵ*.

When the relaxation time is negligible (1/*ϵ* → 0), biodiversity responds instantaneously to habitat conversion (B = *S*(H)) and the ecological feedback on agricultural production is instantaneous. In this case, Fig.2.A shows that transient trajectories converge monotonically to the viable equilibrium (*H*^*^, *B*^*^).

**Figure 2:**
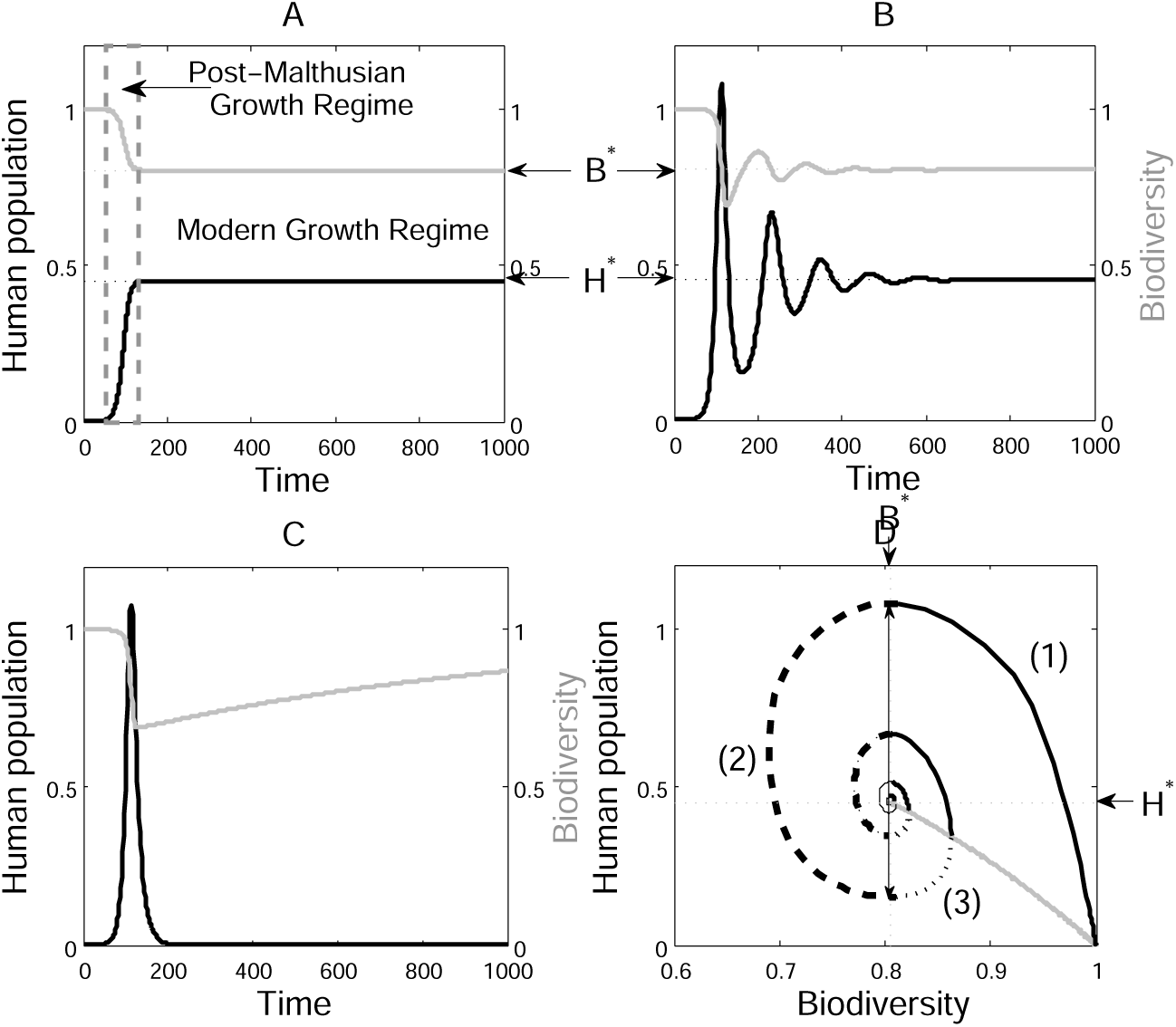
Transient human population and biodiversity dynamics with varying relaxation coefficients (*ϵ*). Parameters are given in Table 1, with *T_m_ =* 1. A: negligible relaxation time (*ϵ* = 10); B: high relaxation time (*ϵ* = 0.02); C: differential relaxation times for biodiversity loss (*ϵ* ^−^ = 0.02) and recovery (*ϵ* ^+^ = *ϵ* ^−^/20); D: phase plane trajectories for biodiversity and human population. Grey curve: negligible relaxation time (*ϵ* = 10); black curve: high relaxation time (*ϵ* = 0.02) leading to the repetition of phases 1 (thick curve), 2 (dashed curve) and 3 (dotted curve) until the equilibrium (*B^*^, H*^**\***^) is reached; double-arrow: maximum amplitude of the transient crisis.

Introducing a time-delay between the dynamics of humans and biodiversity by decreasing ϵ leads to damped oscillations during the transient dynamics to the equilibrium (Fig.2.B). Moreover, if the relaxation rates of species extinction and recovery are distinct (e.g. recovery takes longer than extinction), these damped oscillations can lead to the collapse of the human population (Fig.2.C).

These damped oscillations result from the repetition of three successive phases (Fig.2.D):

(1) **Human population growth and biodiversity decline** (*μ* ≥ 0). During the initial growth phase, the human population reaches its equilibrium size *H*^*^. But at this point, the biodiversity debt is not paid off yet, i.e. biodiversity and ecosystem services are still in excess (B > *B*^*^). As a consequence, the *per capita* consumption is not at equilibrium 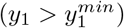, and human population keeps growing until 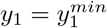.
(2) **Environmental crisis and population decline** (μ ≤ 0). When biodiversity eventually reaches its equilibrium value (B = *B*^*^), both human population growth and habitat conversion stop 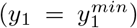. However, the human population now exceeds its equilibrium size (*H* > *H*^*^). This population overshoot results in an over-conversion of natural habitat which generates an additional biodiversity debt. While this debt is paid off, human growth rate becomes negative 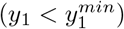 and human population declines.
(3) **Human population and biodiversity recovery** (μ ≥ 0). As the human population declines, the converted surface decreases and allows for the recovery of biodiversity. When biodiversity is not in deficit anymore (B = B*), the human population can reach its equilibrium size, *H*^*^.

### 3.2. Sustainability sensitivity analysis

#### 3.2.1. Sustainability conditions

Transient environmental crises are undesirable from a sustainability perspective, since these collapses result from a decrease in *per capita* agricultural consumption, and thus in human utility (eq. (3)) - which we use as a proxy for human well-being. Let us define an SES as sustainable when its transient dynamics monotonously converge to the long-term equilibrium (i.e. no crisis) and the system experiences a net growth in human utility. Condition (12) ensures that technological change is sufficient to compensate for the effect of biodiversity loss on human utility:

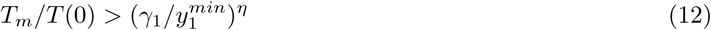

where *T*(0) is the initial technological efficiency. When condition (12) is met, human utility at equilibrium is higher than the initial human utility (*U*^*^ *> U*(0)), but environmental crises may still lead to a decrease in human utility and population size during the transient dynamics. Using dynamical system properties (see Appendix A.3.2), we derive a general condition for the absence of environmental crises:

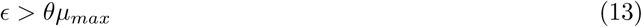

where *θ* is explicitly defined as a function of the economic, demographic, technological and ecological parameters of the SES in Table 1. Condition (13) implies that a given SES is not vulnerable to environmental crises if its ecological dynamics (*ϵ*) is fast enough compared to the growth of its human population (≈ *μ_max_*), for a given set of parameters (*θ*).

Let us rewrite condition (13) as a sustainability criterion, ∆ = *ϵ* − *θμ_max_*. From the expression of *θ* in Table 1, it is straightforward to deduce the effect of some parameters on ∆. In particular, ∆ decreases with the preference for agricultural goods, ∆, since it increases the global demand for agricultural goods. Δ increases with the land operating cost κ, since high operating costs reduce the incentive to convert natural habitat (see Appendix A.4 for more details). The strength of the demographic transition (*b*_2_) also increases the sustainability criterion ∆, by reducing the net growth rate of the human population.

In the next sections, we explore the effect of the other parameters through numerical simulations. We use the default parameters of Table 1 such that conditions (11) and (12) are met, i.e. the SES can reach its desirable equilibrium, and human well-being at equilibrium is higher than the initial well-being. A given SES is sustainable (unsustainable) if its parameters satisfy ∆ > 0 (∆ < 0) and condition (12). Any increase (decrease) in Δ has a positive (negative) effect on long-term sustainability. We then assess the effect of some characteristics of SESs on their steady states (*H*^*^, *B*^*^) and sustainability (∆) by varying a single parameter at a time: the maximum technological efficiency (*T_m_*), the agricultural labor intensity (*α*_1_) and the concavity of the BES relationship (Ω). For each of these parameters, we also plot the effect of the strength of the demographic transition *b*_2_.

#### 3.2.2. Effect of technological change

In our model, production efficiency increases with technological change until a maximum technological efficiency *T_m_.* Increasing *T_m_* results in larger human population sizes and less biodiversity at equilibrium (Fig. 3.A). When the demographic transition is weak (*b*_2_ = 0.2), the effect of technological change on sustainability is negative (Fig. 3.D), and the amplitude of the transient crises rises with technological change (Fig. 3.A and D). On the other hand, technological change has a positive effect on *per capita* and total human utility at equilibrium (Fig. 3.G), since it increases industrial consumption (*y*_2_) and the size of the human population at equilibrium *H^*^*

However, for a stronger demographic transition (*b*_2_ = 2), the effect of technological change on sustainability switches from negative to positive at *T_m_* = 4 (dashed curve in Fig. 3.D). Further increasing *T_m_* allows the industrial consumption *y*_2_ to reduce the net human growth rate to the point where the system does not experience environmental crises (∆ > 0 for *T_m_* > 10). Note that *b*_2_ only affects the transient dynamics of the system, and thus does not prevent high levels of biodiversity loss when technological efficiency rises.

**Figure 3:**
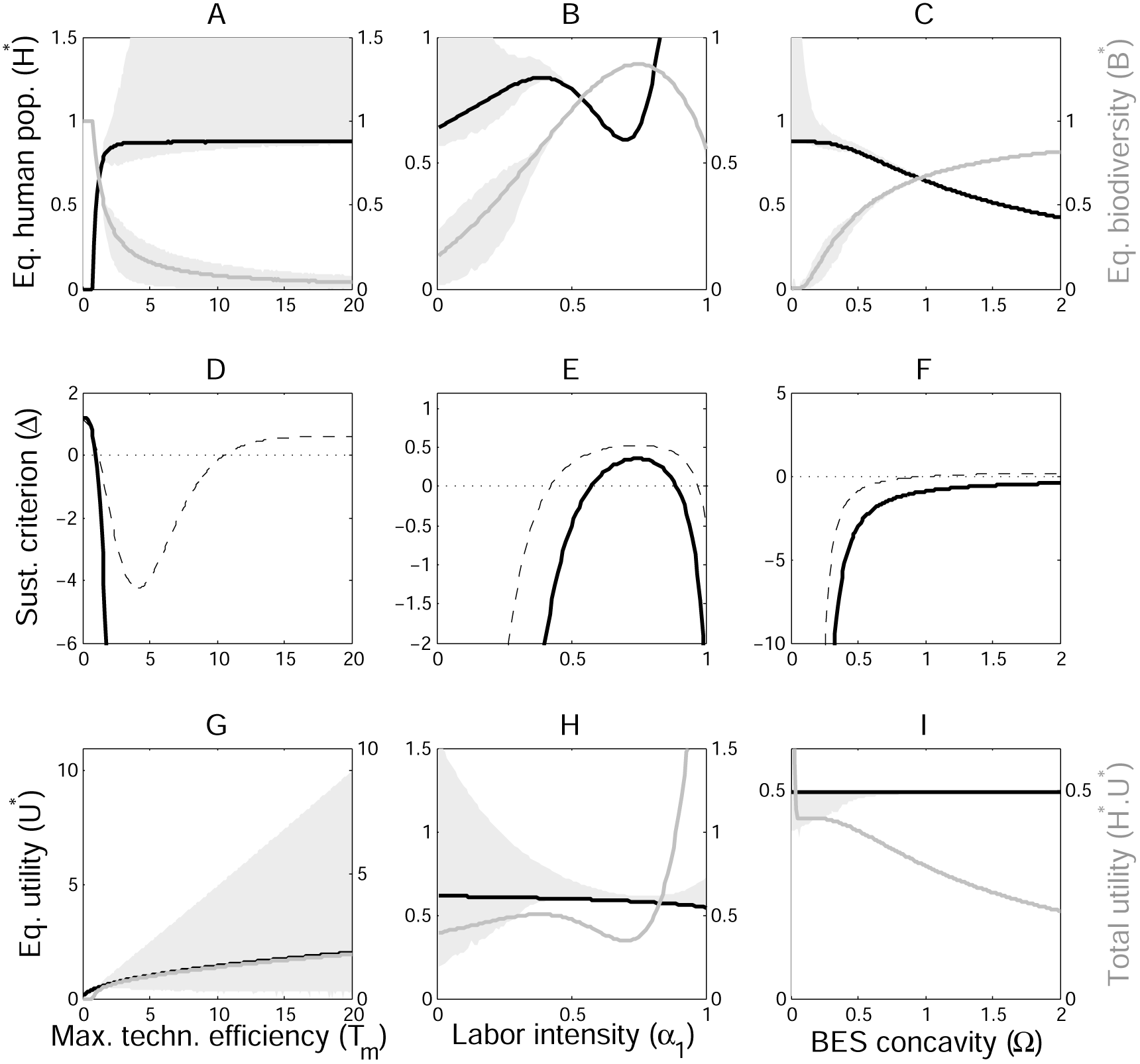
Effect of varying parameter values on the steady states, the sustainability criterion and human wellbeing at equilibrium. Parameters are given in Table 1, with *T_m_* = 1.8 (B-E-H) and *T_m_* = 1.2 (C-F-I). **A** to **C**: Effect of *T_m_, α*_1_ and Ω on the human population size (black curve) and biodiversity (grey curve) at equilibrium, when demographic transition is weak (*b*_2_ = 0.2). Grey areas represent the maximum amplitude of the transient crises as defined in Fig.2.D. **D** to **F**: Effect of *T_m_, α*_1_ and Ω on the sustainability criterion (∆ = ϵ − *θμ_max_*), when demographic transition is weak (solid curve, *b*_2_ = 0.2) and strong (dashed curve, *b*_2_ = 2). Parameter values for which ∆ is positive correspond to sustainable trajectories. **G** to **I**: Effect of *T_m_, α*_1_ and Ω on *per capita* human well-being (*U*^*^, black curve) and total human well-being (*H^*^.U*^*^, grey curve) at equilibrium, when demographic transition is weak (*b*^2^ = 0.2). Grey areas represent the maximum amplitude of the transient variations in well-being.

#### 3.2.3. Effect of labor intensity

As labor intensity *α*_1_ captures the relative use of labor compared to land in the agricultural sector, a moderate increase in labor intensity (*α* _1_ < 0.7) reduces land conversion requirements and benefits both biodiversity and sustainability (Fig.3.B and E). However, further increasing labor intensity (*α* _1_ > 0.7) increases agricultural productivity to the point where it lowers agricultural prices and increases the demand for natural habitat conversion. This rebound effect thus allows the human population and total human utility to rise again, while biodiversity, sustainability and *per capita* human utility decrease (Fig.3.B, E and H). A stronger demographic transition positively affects the sustainability criterion for all labor intensities, and thus partially mitigates the negative rebound effect (dashed curve, Fig.3.E). Note that the shape of the relationship between sustainability and labor intensity depends on the land operating cost, κ. If κ = 1, the bell-shaped curve is centered on *α*_1_ = 0.5. If κ > 1 (κ < 1, resp.), the sustainability criterion is maximum at *α*_1_ *<* 0.5 (*α* _1_ > 0.5, resp.). Indeed, higher operating costs reduce the incentive to convert natural habitat and increase sustainability (Appendix A.4), thus reducing the sustainability-optimal labor intensity.

#### 3.2.4. Effect of the concavity of ecological relationships

The concavity of the ecological relationships, i.e. the SAR and BES relationships, is captured by their parameters *z* and Ω, respectively. Since both parameters have similar effects on ∆, we only present the results for Ω. Commonly observed concave-down BES relationships (Ω < 1) lead to highly populated and biodiversity-poor SESs at equilibrium (Fig.3.C). These SESs are characterized by a higher total human utility (Fig.3.I) and a lower sustainability (Fig.3.F) than SESs with concave-up BES relationships. A stronger demographic transition increases sustainability for both concave-down and -up BES relationships (dashed curve, Fig.3.F).

Modern SESs may thus be particularly vulnerable to the current rise in technological efficiency and reduction in labor intensity - unless the demographic transition is strong enough to counteract their negative effects on long-term sustainability. Sustainable SESs are characterized by higher levels of biodiversity, lower human population sizes, industrial consumption and consumption utility at equilibrium, compared to unsustainable SESs. In section 3.3, we derive sustainability thresholds for land conversion, biodiversity and human population size, and explore their efficiency in preserving the long-term sustainability of SESs through numerical simulations.

### 3.3. Application to land-use management

#### 3.3.1. Integrative sustainability threshold for land conversion

Our sustainability condition ∆ > 0 can be rewritten as *A* < *A_S_* (Appendix A.3.3), where *A_S_* is the sustainable land conversion threshold:

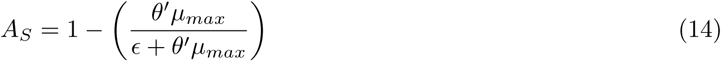

*θ′* is explicitly defined as a function of the economic, technological and ecological parameters of the SES in Table 1. *A_S_* represents the maximum area of natural habitat that a given SES can convert without becoming unsustainable, and depends on the economic and ecological characteristics of the SES. In particular, A*S* decreases as the ecological relaxation rate *ϵ* decreases. As a result, the larger the ecological relaxation time (i.e. the lower *ϵ*), the more natural habitat a SES needs to preserve in order to remain sustainable. Using this land conversion threshold *A_S_*, it is also possible to derive a sustainable biodiversity threshold *B_S_* = (1 − *A_S_*)*^z^*, and a human population threshold *H_S_* = *ϕA_S_*.

#### 3.3.2. Land-use scenarios

In this section, we explore the efficiency of the land conversion threshold (eq.(14)) in preventing environmental crises. To do so, we consider two alternative land-use scenarios: (1) no restriction on land conversion, and (2) conservation of an area of natural habitat 1 − *A_S_*, e.g. through the creation of a protected area. In order to stop land conversion at the sustainable threshold *A_S_* without (directly) limiting the dynamics of the human population, we have to define converted land A as a fourth dynamical variable:

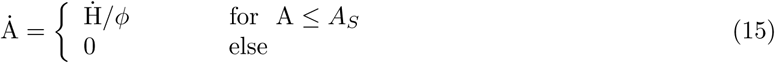

Fig.4 shows that, in an unsustainable SES (i.e. ∆ < 0), the first scenario generates transient environmental crises (Fig.4.A), while the second scenario prevents environmental crises from occurring (Fig.4.B). A precautionary approach to natural habitat conservation appears necessary to prevent biodiversity and human population from exceeding their own sustainable thresholds, thus resulting in environmental crises in the long run.

**Figure 4:**
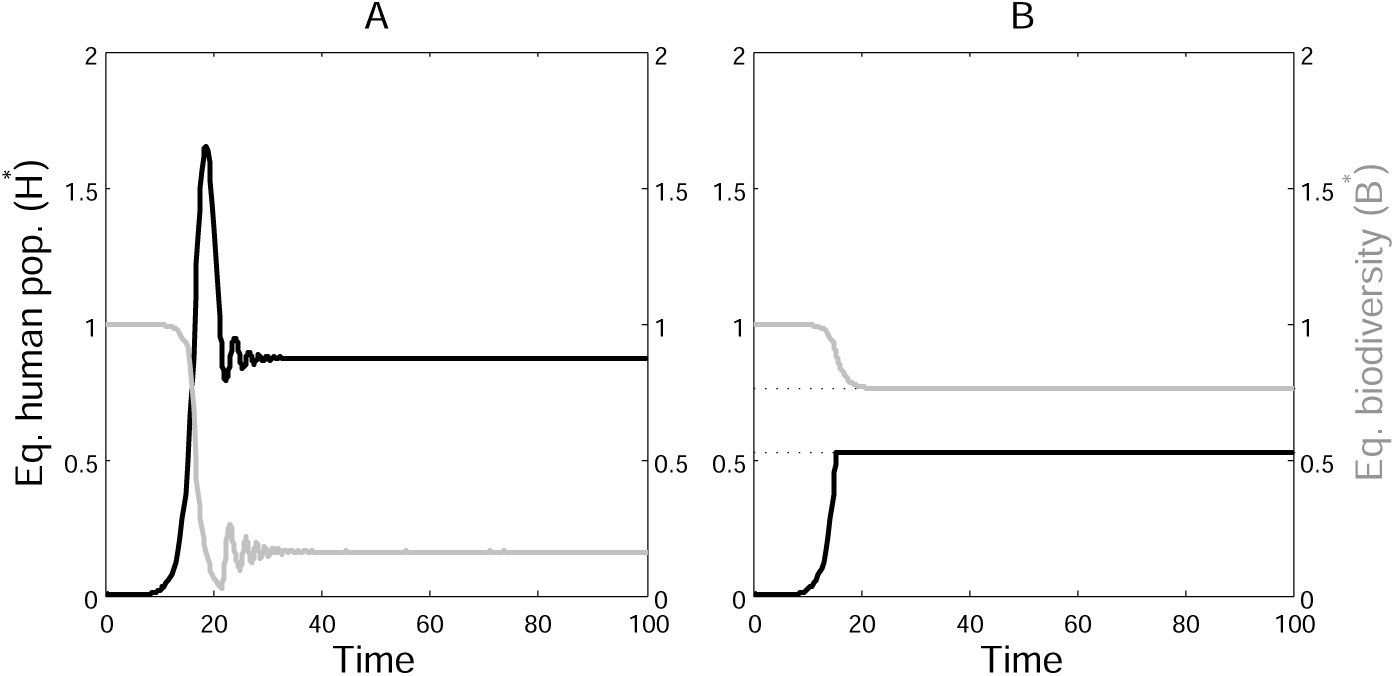
Land-use scenarios. For both scenarios, parameters are given in Table 1, with *T_m_* ***=*** 5. **A: No restriction on land conversion**; dynamics of an unsustainable SES (∆ < 0) with no land-use management. **B: Implementation of the sustainable land conversion threshold**; dynamics of the same SES (∆ < 0) when the sustainable threshold of natural habitat 1 − *A_S_* is set aside.

## 4. Conclusions

We explore the long-term dynamics of coupled SESs by modelling the main reciprocal feedbacks between human population and biodiversity, and show that the temporal trajectories of the human population and biodiversity are not necessarily monotonous. In particular, for large ecological relaxation times, we observe the emergence of transient environmental crises, i.e. large reductions in biodiversity, human population size and well-being. More complex population projection models calibrated with data for the world population, industrialization, pollution, food production and non-renewable resources obtained similar “overshoot and collapse” scenarios (Meadows et al., 1972). Recent updates even suggest that the business-as-usual trajectory of our societies may be approaching such a global collapse (Meadows et al., 2004; Turner, 2014). However, neither these population projection models nor neo-classical economic models consider any long-term ecological feedback of biodiversity on agricultural production, which may worsen these crises scenarios.

The present work thus sheds new light upon the importance of accounting for ecological time lags when studying the sustainability of SESs and defining management policies. Accumulating data on extinction debts (Wearn et al., 2012) and on the relationship between biodiversity and the provisioning of ecosystem services (O’Connor et al., 2017) make it possible to include such time-delayed ecological feedbacks in more complex simulation models. The ability of our toy-model to perform population projections is limited, since many of our economic and ecological parameters do not vary endogenously. For example, the ecological relaxation rate *ϵ* is known to decrease with habitat loss and fragmentation (Hanski and Ovaskainen, 2002), and population density on converted land *ϕ* rose between 1993 and 2009 (Venter et al., 2016), since the rate of land degradation (9%) was lower than that of the human population and economic growth (resp. +23% and + 153%). We calibrated our model using mean values for these parameters during the period 1960-2010, and chose other parameters in order to reproduce the observed human population and TFP growth (Appendix A.5). Population projections suggest strong effects of ecological time lags after 2100, which are worsened if population density keeps rising. These results urge for the inclusion of long-term ecological feedbacks in simulation models of population projections, as is already done in models that couple the economy and climate (Nordhaus, 1993, 2010).

Although detailed policy-relevant population scenarios would require a simulation model of greater complexity, our toy-model allows assessing the qualitative effects of some parameters of SESs on their long-term sustainability. It provides us with an integrative sustainability criterion that captures the vulnerability of an SES to environmental crises, given its economic, demographic, technological and ecological characteristics. Using this criterion, we show that some of the characteristics common to modern SESs, such as high technological efficiency and low labor intensity, are detrimental to their long-term sustainability. Even though land-use efficiency reduces land conversion in the short term, it also increases the time lag between the dynamics of humans and biodiversity and the vulnerability of the SES to environmental crises. Indeed, a more efficient use of land resources releases ecological checks on human population growth, which eventually increases both land conversion and the extinction debt. Moreover, the commonly observed concave-down SARs and BES relationships exacerbate this effect, since a given amount of habitat conversion translates into a lower loss of biodiversity and ecosystem services with concave-down than with concave-up relationships.

We show that counteracting forces such as the demographic transition and natural habitat conservation can mitigate environmental crises - provided that the ecological relaxation rates and the sensitivity of the human growth rate to industrial consumption are high enough. Indeed, the demographic transition is often presented as the solution to accelerate economic development and reduce environmental impact in developing countries (Bongaarts, 2009; Brander, 2007). However, recent projections cast doubt upon its stabilizing potential (Bradshaw and Brook, 2014). Our results also show that, in order to efficiently counteract time-delayed biodiversity feedbacks, natural habitat conservation should take place when biodiversity is still abundant. Alternatively, habitat restoration can also help mitigating crises, and century-long extinction debts may be seen as windows of conservation opportunity (Wearn et al., 2012). However, our results suggest that we should not rely on habitat restoration, since restoration delays are often longer than extinction debts (Naaf and Kolk, 2015; Maron et al., 2012), which worsens environmental crises.

The lack of such considerations about ecological time lags in current conservation policies calls for a precautionary approach to natural habitat conservation. To this end, we provide an integrative land conversion threshold that captures the long-term ecological dynamics of species confronted with the destruction of their habitat, given the economic, technological and demographic parameters of an SES. The smaller the ecological relaxation rate (*ϵ* → 0), the more natural habitat should be preserved in order to avoid environmental crises. Since relaxation rates decrease with habitat loss and fragmentation (Hanski and Ovaskainen, 2002), the sustainable amount of natural habitat may increase when more and more natural habitat is lost.

Regarding rates of current biodiversity loss (Newbold et al., 2016), it is thus crucial to determine how much natural habitat we need to preserve in the long run. Recent attempts to define global conservation thresholds for biodiversity and land use (Steffen et al., 2015) neglect the interaction between the social and ecological components of SESs. For instance, as biodiversity moves closer to its own potential threshold, it reduces the land-use change threshold (Mace et al., 2014). There is also a lack of context-dependency for the application of these thresholds at local scales, as SESs with different sets of economic, technological and ecological characteristics present different thresholds. Our theoretical model of a coupled SES proposes a way to move beyond these limitations. Our integrative thresholds for land conversion and biodiversity loss are not defined independently from each other, but related through the system’s parameters. As a consequence, it is possible to predict how a change in one of the thresholds affects the other, and to define context-dependent land-use policies that prevent environmental crises from occurring.

Despite its simplicity, our model may be seen either as a thought experiment about the sustainability of our planet, or as a representation of a local SES - which could be connected to other SESs by considering trade, human migrations and species dispersal. Improvements to the model include adding property rights and trade in order to gain realism in land-use change dynamics (Lambin et al., 2001), and internalizing the value of ecosystem services via payments for ecosystem services and taxes on land conversion (Costanza et al., 2014). Finally, our results emphasize the critical need for a better assessment of ecological time lags and how they feed back on human systems. Reducing uncertainties on ecological time lags effects is crucial to be able to define integrative thresholds and foster sustainability of our planet, which may be seen as a global social-ecological system (Ehrlich and Ehrlich, 2013). The pursuit of such objectives requires taking into account the main internal feedbacks and specificities of coupled social-ecological systems, which dynamical system modelling allows.

## Acknowledgements

This work was supported by the TULIP Laboratory of Excellence (ANR-10-LABX-41) and the Midi-Pyrénées Region. We would like to thank two anonymous reviewers for their careful reading of our manuscript and their insightful comments and suggestions. We also thank François Salanié, Claire de Mazancourt, Bart Haegeman, Audrey Valls, Robin Delsol, David Shanafelt, Charles Perrings and Ann Kinzig for valuable discussions and helpful comments on earlier versions of the manuscript.

## A. Appendices

### A.1. Endogenous or Exponential Technological Progress

Technological efficiency T represents the Total Factor Productivity (TFP) of the industrial and agricultural sectors. Following recent concerns regarding the potential for continued TFP growth in the future (Gordon, 2012; Shackleton, 2013), we choose to model technological change as a logistic function, such that technological efficiency grows at an exogenous rate *σ,* until it reaches a maximum technological efficiency (*T_m_*).

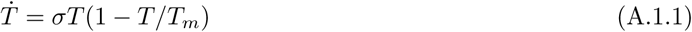

This function allows us to retain analytical tractability, and for instance, derive analytical sustainability criteria. We could also have modeled technological change as endogenous, e.g. driven by the rate of human population growth :

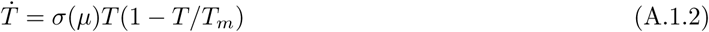

where *σ*(*μ*) is a function of the human growth rate, for instance *σ*(*μ*) = *μ* (*B, T*). The behaviour of the system is not qualitatively affected by this assumption. However, the system becomes analytically intractable.

Another possibility is to assume an exponential technological progress, i.e. technological efficiency keeps growing instead of reaching a maximum technological efficiency *T_m_*:

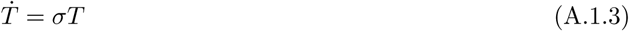

In this case, the system has no global equilibrium (*H^*^,T^*^,B^*^*) and no analytical results. However, as technological change increases the strength of the demographic transition, it reduces the growth rate of the human population (*μ*(*T, B*) → 0) along with land conversion, so that the human population and biodiversity reach a stationary state in the long run.

In both cases (endogenous or exponential technological change), however, our results would not be qualitatively affected. Indeed, with endogenous technological change and a strong demographic transition (i.e. high value of the sensitivity of the human growth rate to industrial consumption *b*_2_), the effect of the maximum technological efficiency *T_m_* on the vulnerability to environmental crises switches from negative to positive and reduces the amplitude of the transient environmental crises (Fig. A.5.A). This corresponds to the dashed curve in fig.3.D (*b*_2_ = 2). Similarly, with an exponential technological change, the effect of varying labor intensity *α*_1_ is non-linear (Fig. A.5.B), as it first reduces environmental crises before increasing human population size again, through an economic rebound effect. This corresponds to fig.3.B and E.

**Figure A.5:**
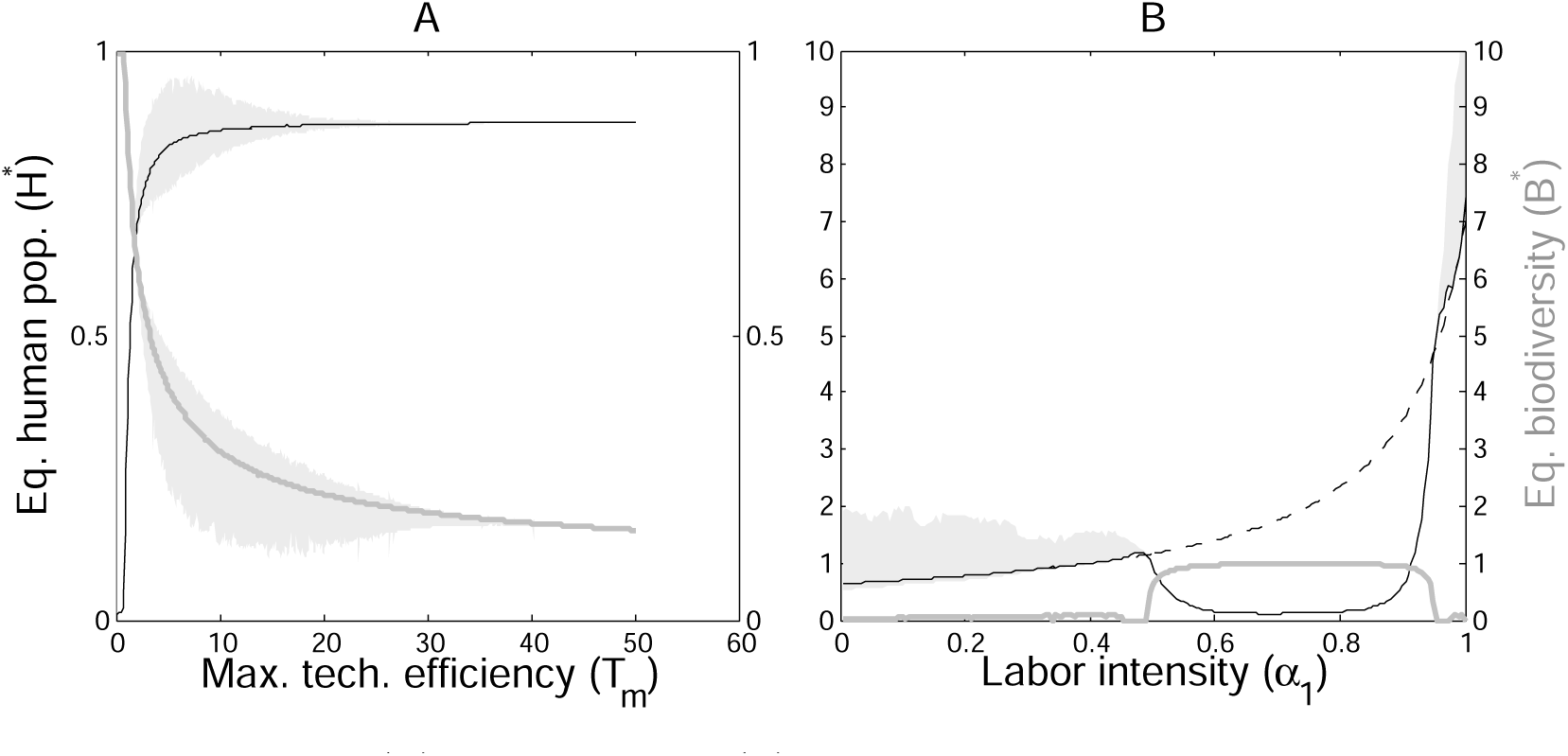
Effect of endogenous (A) and exponential (B) technological change on long-term sustainability. **A:** effect of an endogenous technological change (*σ* = *μ* (*B, T*) on the transient and long-term dynamics of the system, when the maximum technological efficiency *T_m_* varies. Grey areas represent the amplitude of the transient environmental crises. Parameters are given in Table 1, with Ω = 2 and *b*_2_ = 0.5.; B: effect of an infinite technological change (Ṫ = *σ)* on the transient and long-term dynamics of the system, when the labor intensity *α*_1_ varies. Parameters are given in Table 1, with Ω = 2, *b*_2_ = 0.1 and *σ* **=** 0.1.

### A.2. Economic derivations

In this section, the aim is to derive the *per capita* agricultural and industrial consumptions at the market equilibrium, i.e. when the production supply equals the demand of the human population. The human population is assumed to be composed of identical agents, with preferences *η.* Each agent allocates part of his revenue *w* to buy agricultural goods, and the rest to buy industrial goods, which generates an aggregate demand for the human population H (section A.2.1). On the supply side, firms produce agricultural and industrial goods given the costs of production inputs, i.e. land and labor, which gives the aggregate supply of the production sectors (section A.2.2). Section A.2.3 solves for the general equilibrium model, and derives the *per capita* consumptions at the market equilibrium. Section A.2.4 then derives the rate of land conversion at the economic market equilibrium.

#### A.2.1. Consumer Optimization

Overall, each agent supplies one unit of labor, so that his revenue is the wage w. Let *U*(*y*_1_,*y*_2_) be the individual consumer well-being, where *y*_1_ and *y*_2_ are *per capita* consumption rates of agricultural and industrial goods. Agents maximize their well-being 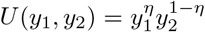 under the revenue constraint *p*_1_*y*_1_ + *p*_2_*y*_2_ *≤ w,* where *p*_1_ and *p*_2_ are the prices of agricultural and industrial goods respectively, w the *per capita* income, and *η* the preference for agricultural goods. To solve this maximization problem, we define the Lagrangian:

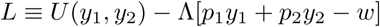

First order conditions are:

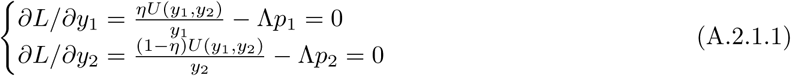

Adding both conditions yields *U*(*y*_1_,*y*_2_)/Λ = *p*_1_*y*_1_ + *p*_2_*y*_2_ = *w*, and solving (A.2.1.1) for *y*_1_ and *y*_2_, and substituting for *U*(*y*_1_,*y*_2_)/Λ yields the aggregate demands:

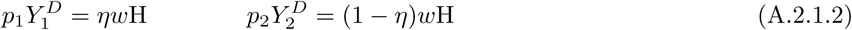

#### A.2.2. Firms Optimization

Firms in sectors 1 and 2 produce a quantity Y_i_ (*i* = 1, 2) of output, using labor *L_i_*, and land *A_i_*. A units of land and H units of labor are available. Agricultural firms occupy *A_1_* units of land, and industrial firms occupy *A*_2_ units of land, such that *A* = *A*_1_ + *A*_2_. This comes at an operating cost (including clearance and maintenance) of *κ* units of labor per unit of land, so that each firm maximizes a profit Π_*i*_ = *p_i_Y_i_* − *wL_i_* −*κwA_i_*. The production functions *Y_i_* are:

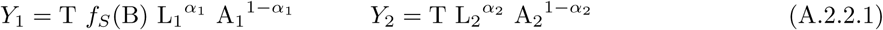

where *f_S_* (B) is the ecological feedback, T captures production efficiency and *α_i_*, is labor intensity in sector *i*. Therefore, first order conditions are:

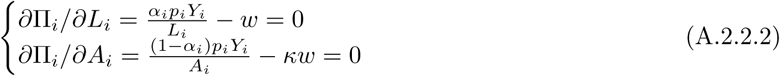

Adding both lines of (A.2.2.2) gives the total supply in sector *i*:

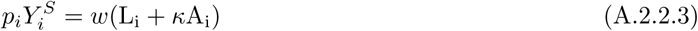

#### A.2.3. Economic General Equilibrium

Markets’ clearing for the agricultural and industrial sectors 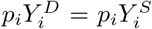 (eq. (A.2.1.2) and (A.2.2.3)) yields:

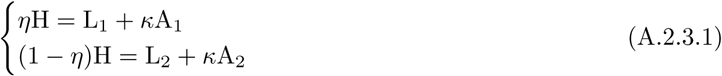

Input factors in each sector verify *L_i_*/*α_i_* = κ*A_i_*/(1 − *α_i_)* (see eq. (A.2.2.2)), so that the equilibrium land allocations are:

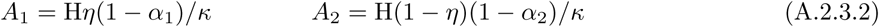

and the equilibrium labor allocations are:

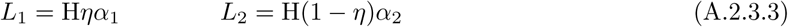

#### A.2.4. Land Conversion

The allocation of total labor H between agricultural production (*L*_1_), industrial production (*L*_2_) and land conversion (κA) writes *L*_1_ + *L*_2_ + *κ*A = H. Replacing *L*_1_ and *L*_2_ by their optimal allocations (eq.(A.2.3.3)), we deduce the relationship A = *H/ϕ* between the human population size H and the converted land A, where the density of the human population per unit of converted land is:

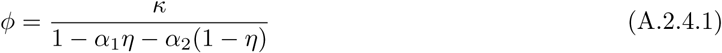

#### A.2.5. Per capita consumptions

Using the equilibrium allocations of labor *L_i_* and land *A_i_*, the *per capita* agricultural and industrial consumptions *Y/H* (eq.(A.2.2.1)) can be rewritten:

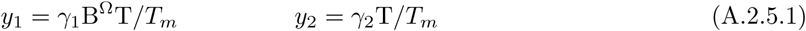

where

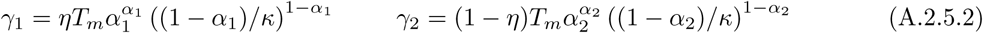

### A.3. Dynamical System Analysis

#### A.3.1. Equilibria and Stability Condition

The Jacobian matrix at the viable equilibrium (*H*^*^, *T_m_*, *B^*^*) writes:

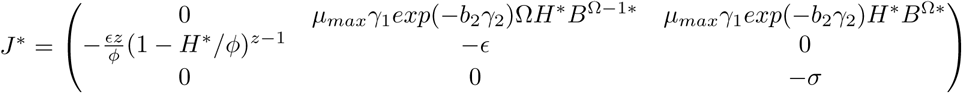

The eigenvalues λ*_i_* (*i* =1:3) of the system are solutions of *Det*(*J^*^ −* λ*I*) = 0, where

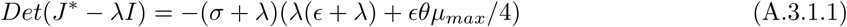

and

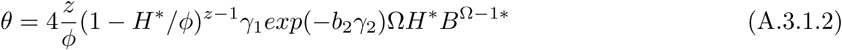

The first eigenvalue is λ_1_ = *−σ,* and the two others are solution of the characteristic equation λ^2^ + *ϵ*λ + *ϵθμ_max_/4* = 0, which discriminant is:

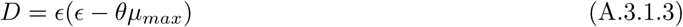

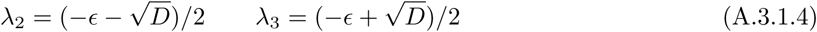

The viable equilibrium is stable if all the eigenvalues have negative real parts, i.e. 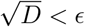, which gives a stability condition for the viable equilibrium (H^*^*,T_m_,* B^*^):

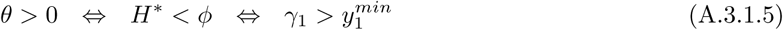

#### A.3.2. Sustainability Condition

For parameters such that the viability condition 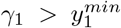 is met, there are two possible transient dynamics depending on the eigenvalues of the system: (1) for real eigenvalues, a monotonous convergence to equilibrium, and (2) for complex conjugate eigenvalues, damped oscillations. The former case stands for a sustainable system, while the later one stands for an environmental crisis. The eigenvalues are real if and only if the discriminant of the characteristic equation is positive, so that a sustainability condition for our system is:

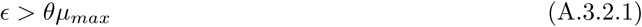

A sustainability criterion for our system is ∆ = ϵ − *θμ_max_,* where ∆ > 0 stands for sustainable trajectories.

#### A.3.3. Sustainability Thresholds

Using the relationship between the biodiversity and human population size at equilibrium *H^*^* = *ϕA^*^, θ* (eq.(A.3.1.2)) can be rewritten as a function of the equilibrium converted area *A*^*^. The sustainability criterion ∆ then rewrites:

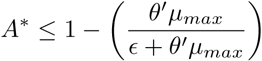

where 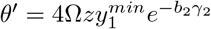, and *ϵ* is the ecological relaxation rate. Over a sustainable trajectory (∆ > 0), converted land never exceeds its equilibrium level, i.e. A ≤ A^*^. Condition (A.3.3) is thus equivalent to A ≤ *A*_S_, where *A_S_* is the sustainable land conversion threshold of our system. Similarly, the sustainability criterion can be written as a human or a biodiversity threshold as follows:

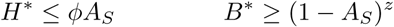

### A.4. Effect of Land Operating Costs on Sustainability

The exploitation of converted land comes with operating costs at each period of time, which include the initial cost of natural habitat conversion, and the cost of land maintenance. Operating costs *κ* are in units of human labor per unit of converted land. High operating costs are beneficial to the long-term sustainability of the SES (Fig.A.6.B), since they reduce the incentive to convert natural habitat - in a similar manner to taxes on converted land. Therefore, biodiversity at equilibrium increases with *κ* (Fig.A.6.A) while consumption utility decreases (Fig.A.6.C). The effect of *κ* on the size of the human population at equilibrium *H*^*^ is non-linear (Fig.A.6.A), since low values reduce the population density *ϕ*, and high values make the system unviable 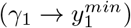, i.e. natural habitat conversion becomes to expensive.

**Figure A.6:**
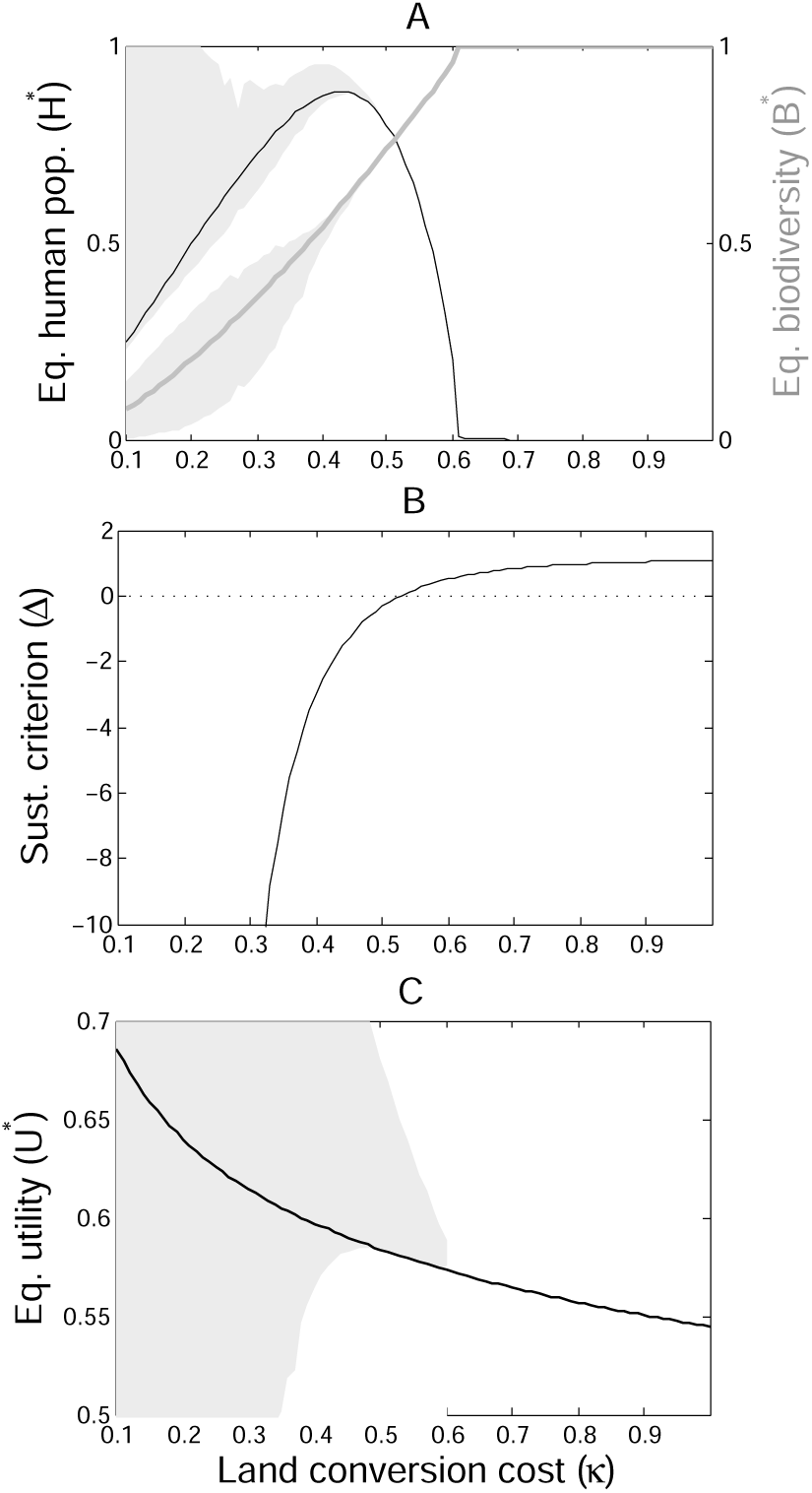
Effect of the land conversion cost (*κ*). Effect of varying *κ* on the viable equilibria (*H^*^,B^*^*), with default parameters of Table 1 and *T_m_* = 1.8. The grey area represents the amplitude of the transient crises. **B**: Effect of κ on the sustainability criterion (∆). Values of *κ* for which ∆ is positive correspond to sustainable trajectories. **C**: Effect of *κ* on human well-being at equilibrium. The grey area represents the amplitude of the transient variations in well-being.

### A.5. Model Calibration

The model was calibrated on the period 1960-2010, using estimates for some parameters, e.g. *z* = 0.27 (Wearn et al., 2012), and choosing aggregate and demographic parameters in order to reproduce the observed changes in human growth rate (≈ 2% in 1960, ≈ 1.2% in 2010), human population size (3 billions in 1960, 6.9 billions in 2010) and TFP growth (nearly tripled) between 1960 and 2010. Trajectories lead to a population size around 15 billions people in 2100, slightly higher than the medium fertility predictions of the United Nations (11.2 billions).

In order to explore the effect of relaxation rates on the population projections, we choose two relaxation rates corresponding to half-lives of 23 and 35 years. Top panels of Fig. A.7 show that trajectories are unaffected by ecological time-lags between 1960 and 2100, but diverge greatly afterwards. Note that the halflives chosen are much shorter than the usual estimate of 57 years in local communities (Wearn et al., 2012). The longer the half-life, the larger the growth of the human population and the resulting crisis, since the long-term equilibrium is unchanged (with constant parameters, and especially constant population density).

**Figure A.7:**
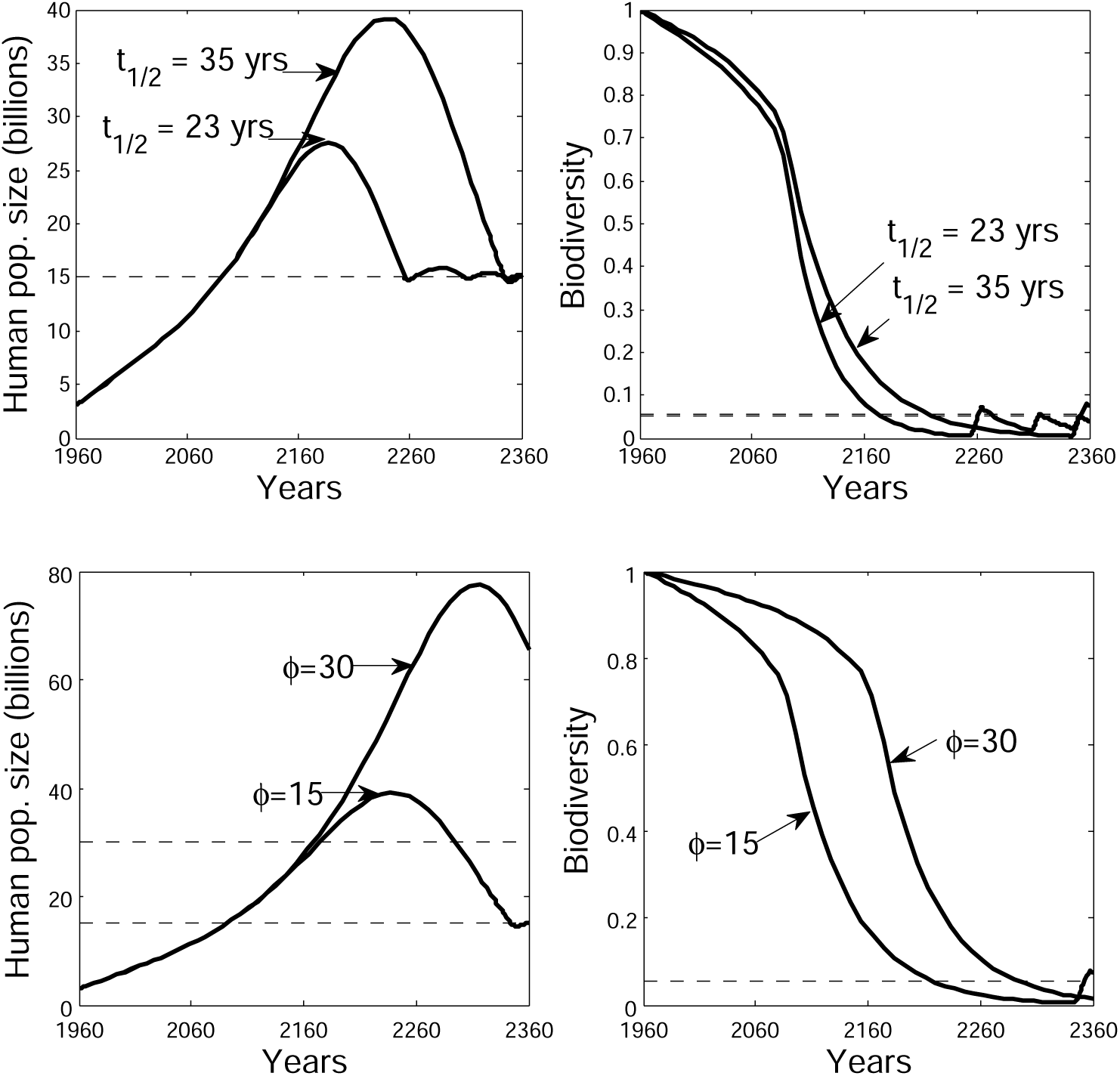
Model calibration and population projections. **Top panels:** Projected population and biodiversity trajectories for varying ecological relaxation rates (*ϵ*), with *ϕ* =15 corresponding to the mean population density between 1960 and 2010. Half-lives *t*_1/2_ = *ln*(2)/*ϵ* measure the time it takes to loose half of the species that are doomed to extinction. A half-life of 35 years corresponds to a relaxation rate *ϵ* = 0.02, and a half-life of 23 years corresponds to a relaxation rate *ϵ* = 0.03. Bottom panels: Projected population and biodiversity trajectories for varying human population densities (*ϕ*), with *ϵ* = 0.02. *ϕ* = 15 corresponds to the mean density of human population on total deforested land between 1960 and 2010. Parameter values: *μ_max_* = 0.05; *b*_2_ = 2.29; *T_rn_* = 0.15; *γ*_1_ = 60; *γ* _2_ = 5; 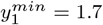; *σ* = 0.05. Dashed lines: long-term equilibria.

Bottom panels show the effect of an increase in population density on the long-term trajectories, with a 35 years half-life. Once again, trajectories are affected by population density only after 2100, the larger density leading to higher human population sizes in the long term. Moreover, larger population densities result in a higher amplitude of the crisis, thus worsening the effect of relaxation rates.

These results suggest that current trends of increasing population density and ecological relaxation times may lead to highly unsustainable pathways in the long run.

